# Interstrand Crosslinks in Donor DNA Boost Gene Editing in Human Cells

**DOI:** 10.1101/2022.03.11.484022

**Authors:** Hannah I. Ghasemi, Julien Bacal, Amanda Yoon, Carmen Cruz, Jonathan Vu, Brooke Gardner, Chris D. Richardson

## Abstract

Co-introduction of targeted nucleases and DNA/RNA templates encoding new genomic sequence is the basis for rapid, effective, and iterable gene editing workflows for therapeutic, agricultural, and basic science applications. Extensive optimization of reagent delivery and nuclease activity have improved genome editing workflows, but comparatively few efforts have been made to alter the gene editing activity of template molecules. Here, we report template DNA modified with interstrand crosslinks (ICLs) – xHDRTs - increases editing frequencies in Cas9-directed gene editing workflows by up to five-fold. xHDRTs increase gene editing frequencies independent of DNA template topology, amount of sequence added, or cell type. Gene editing using xHDRTs requires the DNA repair kinase, ATR, and partially requires Fanconi Anemia proteins, including FANCA, but is independent of other ICL-repair pathways. Covalent modification of donor DNA thus presents a compelling opportunity to improve nonviral gene editing workflows.

## Introduction

CRISPR/Cas9 enables gene editing via DNA double strand break (DSB) generation and subsequent activation of cellular DNA repair pathways. Depending on the repair pathway that is engaged, outcomes can include disruption of the targeted gene or replacement with new sequence that restores or introduces functionality^1^. These latter gene replacement events require the delivery of template DNA encoding new sequences to levels that support gene replacement but do not adversely affect cell viability. In translational applications, template molecules are often delivered by viral vectors. While effective, viral workflows are expensive, difficult to scale, and potentially toxic to cells. Use of non-viral template DNA is thus an appealing alternative, but the efficiency and acute toxicity of non-viral templates can be inferior to viral delivery^2^. Improved non-viral gene editing would be a powerful approach to unravel DNA repair mechanisms, a useful laboratory technique, and a promising strategy for the treatment of a multitude of diseases^3^.

One high efficiency non-viral gene editing strategy co-delivers ribonucleoprotein (RNP) formulations comprising the targeted nuclease Cas9, a single guide RNA (sgRNA), and a template molecule that contains homology to the region being edited as well as the sequence to be modified or inserted^4^. These RNPs introduce DSBs at targeted regions in the genome, which are then repaired by error prone end joining (EJ) processes that rejoin the ends of the break, or homology-directed repair (HDR) processes that resolve DSBs using sequence encoded in a separate template molecule^1^ (**Fig. S1A**). Use of HDR to introduce new DNA sequence into targeted locations enables exciting gain-of-function applications^5^. Strategies to increase HDR frequency may therefore improve outcomes and decrease costs in laboratory and biomedical workflows.

Gains in non-viral HDR efficiency have been achieved through optimization of editing reagents, including protein engineering of Cas9 and related nucleases^6^, improving delivery of reagents into cells^7^, biophysical optimization of RNP parameters^8^, and optimization of size and orientation of the homology region of template DNA^9,10^. Parallel lines of research have focused on defining the cellular response to editing reagents with the goal of redirecting repair events through desired repair pathways^11,12^. These studies have developed key insights into DNA repair processes that underlie gene editing, but with few exceptions^13,14^, it has been hard to translate this understanding into treatments that bias DSB repair towards desirable outcomes. One limitation may be an inability to upregulate DNA repair processes that contribute to DSB repair. For example, we and others demonstrated that non-viral gene editing requires the Fanconi Anemia (FA) pathway and that these FA proteins localize to DSBs^11,15,16^. However, overexpression of key FA genes failed to increase HDR beyond frequencies seen in control strains^11^.

We reasoned that adding substrates for desired DNA repair pathways to template DNA would be an orthogonal approach to activate desired DNA repair activities. Here we report that adding interstrand crosslinks (ICLs) – substrates for the FA DNA repair pathway – to template DNA stimulates HDR by approximately three-fold on a per mole basis in human cell lines, iPS cells, and stimulated T-cells, without increasing off-target mutation frequencies.

## Results

We adapted a nonviral gene editing workflow to measure the effect of covalent modification of double stranded HDR templates (HDRTs) on gene editing efficiency. Interstrand crosslinks (ICLs) added to an HDRT – which we refer to as xHDRTs – dramatically improve editing rates in non-viral gene editing workflows (**Fig. 1A**). ICLs are perturbing, cytotoxic DNA lesions, which covalently tether both DNA strands together, and are repaired in human cells by replication- and transcription-coupled mechanisms^17–19^. Common crosslinking agents include psoralen, which crosslinks TA sites^20^, and cisplatin, which crosslinks GC sites^21^. Both psoralen and cisplatin crosslinking reagents stimulate HDR when used to make xHDRTs, suggesting that the HDR stimulation is general to ICLs and not to a specific chemistry (**Figs 1A and S1B**). Psoralen crosslinking requires long-wave UV irradiation, thus unreacted psoralen cannot form ICLs in cells (where no UV exposure occurs), so we prioritized the development of psoralen-derived xHDRTs. Incubation of HDRTs with varying concentrations of psoralen and 365nm UV radiation created xHDRTs that increase integration of GFP into the HBB locus of human cells ∼three-fold (**Fig. 1A**). Given the magnitude and counter intuitive nature of this effect, we ruled out potential confounding mechanisms. GFP was not expressed from the xHDRT itself, as psoralen ICLs inactivate transcription from reporter genes expressed on the xHDRT (**Fig. S1C**). Nor was this effect caused by nonspecific integration of donor sequence into the genome, as xHDRTs that attach GFP to the N-terminus of LMNB1 produce signal consistent with the fusion protein, and side products indicative of frequent off-target insertion do not appear in the edited samples (**Fig. S1D**). Addition of xHDRTs to cells causes a slight enrichment of cells in the G2 phase of the cell cycle over asynchronous controls, but this is indistinguishable from cells treated with uncrosslinked templates (**Figure S1E**). We note that HDRTs containing primarily thymidine dimers^22^ caused by longwave UV radiation did not support elevated levels of HDR (**Figure 1A, 0 mM**), and so increased editing is specific to ICLs and not nonspecifically caused by damaged donor DNA. Overall, xHDRTs can be used in existing gene editing workflows to boost HDR by approximately three-fold on a per-mole basis.

**Figure 1:**
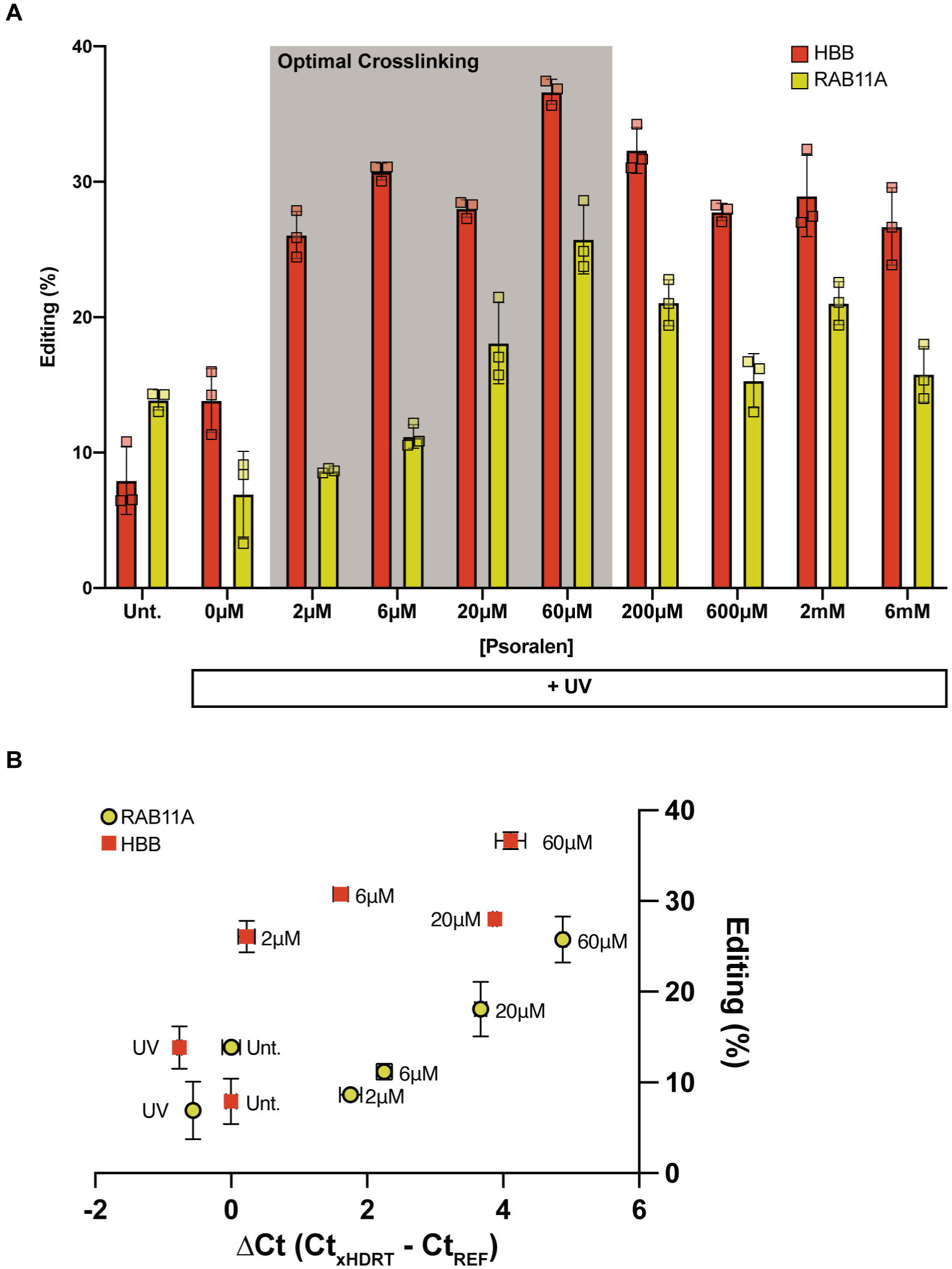
Modification of HDRTs with an optimal number of interstrand crosslinks increases HDR during gene editing. **(A)** Percent of cells GFP positive after editing with pSFFV-GFP (HBB) or N-terminal GFP-fusion (RAB11A) constructs in human K562 myeloleukemia cells. xHDRTs were produced by treatment with the indicated amount of psoralen and UV exposure. Unt – ethanol precipitated donor DNA (no UV, no psoralen). **(B)** Percent of cells GFP positive (y-axis) as a function of qPCR signal loss (x-axis), an approximation of crosslinks per unit length, for xHDRTs produced with the indicated psoralen concentration. Data displayed as the mean ± SD of n=3 biological replicates.

Psoralen crosslink density is a function of the TA content of the DNA, the psoralen concentration, and the UV dosage and may thus vary between HDRTs. To estimate the optimal number of ICLs per xHDRT, we developed a qPCR-based assay that approximates the number of crosslinks within a given DNA molecule (**Fig. S1F**). Using primers that amplify a 94 basepair region of the HDRT plasmid backbone, we determined the probability that at least one crosslink has been introduced in this region. We calculated the ratio (expressed as ΔCt) of qPCR signal produced from xHDRTs generated with different psoralen concentrations or uncrosslinked templates. The editing activity of xHDRTs relative to uncrosslinked controls peaked at three-fold, which occurs at a mean ΔCt value of 4.5 (**Figure 1B**). This translates to a crosslink density of ∼10 crosslinks per kilobase. These parameters were consistent for xHDRTs homologous to the HBB and RAB11A loci.

To define the generalizability of our xHDRTs, we tested these constructs in the context of different donor DNA topologies and sequences. xHDRTs boost gene editing in the context of linear and circular double stranded molecules, and for HDR payloads including three nucleotide SNPs (∼five-fold), single kilobase constructs (∼two-fold), and multi-kilobase constructs (∼three-fold) (**Fig. 2A and S2A**). To validate our approach in other human cell lines, we confirmed that xHDRTs increase HDR of a single kb insertion by ∼two-fold as compared to an uncrosslinked template in additional cell lines, including UMSCC1 and HEK293T cells (**Fig 2B**). We also validated that xHDRTs stimulate HDR in stimulated human T-cells (∼four-fold) (**Fig 2C**) and iPS cells (∼three-fold) (**Fig 2D**), which are useful cell lines for regenerative medicine or immuno-oncology applications. Our overall conclusion is that xHDRTs boost gene editing regardless of payload or target cell type.

**Figure 2:**
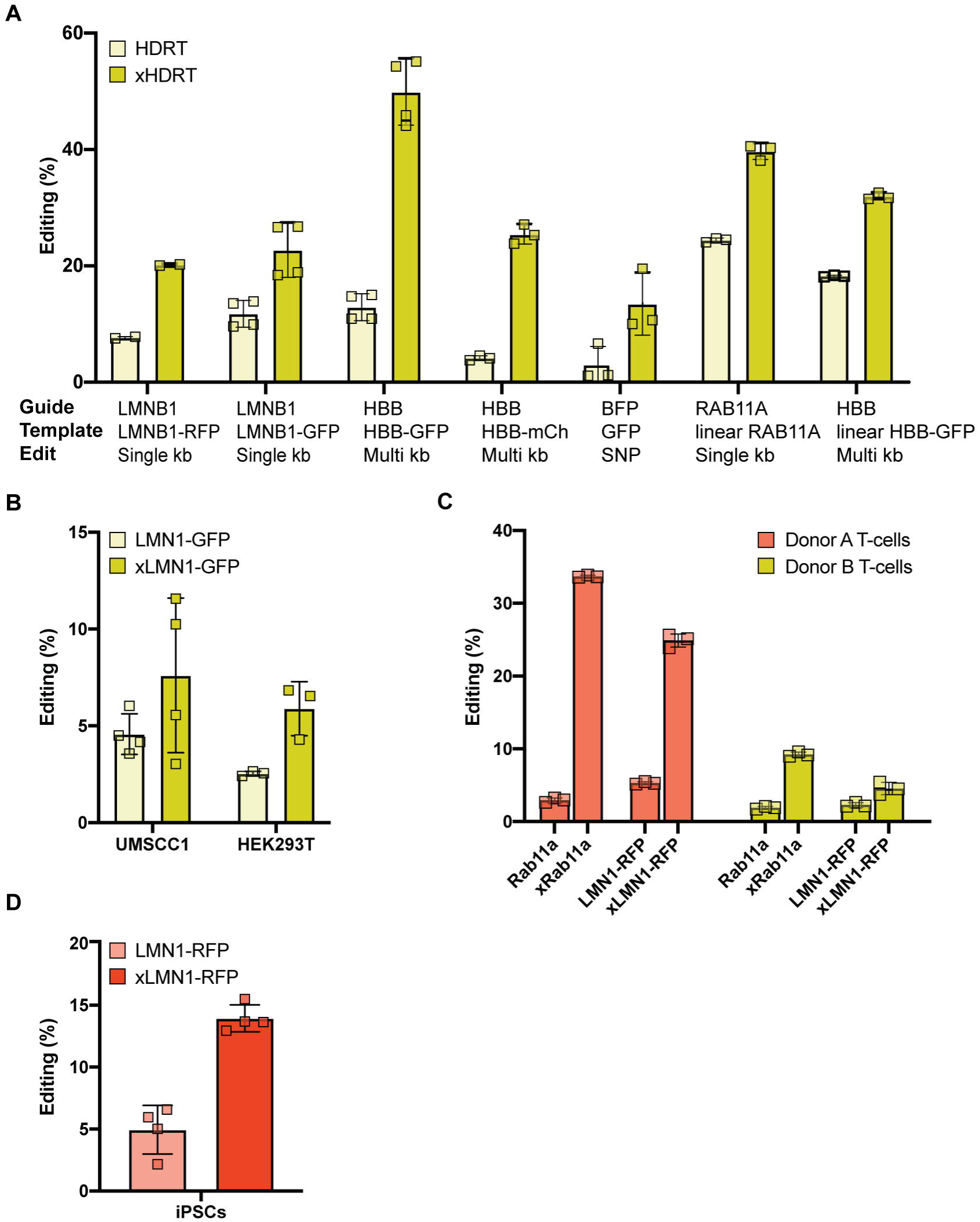
xHDRTs increase HDR in broad gene editing applications. **(A)** Percent incorporation of single kilobase (LMNB1, RAB11A), multi kilobase (HBB), and SNP (BFP) sequences using plasmid (left) or linear (right) double stranded DNA in K562 cells. **(B)** Percent incorporation of a single kilobase sequence at the LMNB1 locus of UMSCC1 and HEK293T cells. **(C)** Percent incorporation of a single-kilobase construct at the LMNB1 or RAB11A loci in primary T-cells isolated from two donors. **(D)** Percent incorporation of a single-kilobase construct at the LMNB1 locus in iPS cells. Data were obtained by flow cytometry and displayed as the mean ± SD of at least n=3 biological replicates.

xHDRTs contain DNA lesions that are potentially mutagenic; however, we see no evidence that HDR using xHDRTs is more mutagenic than HDR using uncrosslinked templates. This is apparent during fluorescent tagging of endogenous genes, where we observe a ∼3-fold increase in GFP cells rather than any decrease caused by frame- or codon-disrupting mutations introduced by ICLs in the GFP donor sequence (**Figure 2**). We further investigated mutation frequencies during SNP editing experiments and observed no increase in cumulative mutation frequencies in a window surrounding the Cas9 cut site relative to those observed during editing with RNP alone or with RNP and uncrosslinked template (**Figure S2B**). However, we note that the background mutation rate (the noise) of our amplicon sequencing data is approximately 2×10^−3^ per nucleotide (**Figure S2B Unedited**), which is many orders of magnitude higher than the fidelity of *in vivo* DNA metabolism, e.g. replicative fidelity (∼1×10^−8^)^23^. To boost the sensitivity of our assay, we focused on TA-sites, which are the substrates for psoralen crosslinks, and are present in the 50bp window surrounding the BFP (2) and HBB (1) cut sites. We observe no increase in mutation rates at these sites in xHDRTs relative to uncrosslinked controls (**Figure S2C**). Overall, we conclude that xHDRTs promote HDR without decreasing HDR fidelity.

xHDRTs could boost HDR through biophysical parameters, e.g. by altering delivery of editing reagents, or by altering the recognition of xHDRTs by cellular DNA repair pathways. To determine whether ICLs are detected in xHDRTs or trigger a cell-wide response that favors HDR, we tested if the ICL had to be present in *cis* on the homologous template molecule. We simultaneously transfected two plasmids, one containing homology to the break site, and one lacking homology, with ICLs present on the homologous, nonhomologous, or neither template DNA. Only ICLs on the homologous template, but not the nonhomologous template, boosted HDR at the LMNB1 and HBB loci (**Fig. 3A**). This experiment indicates that the xHDRT acts through local perturbations on the template DNA that improve HDR efficiency, rather than globally priming the cell for HDR.

**Figure 3:**
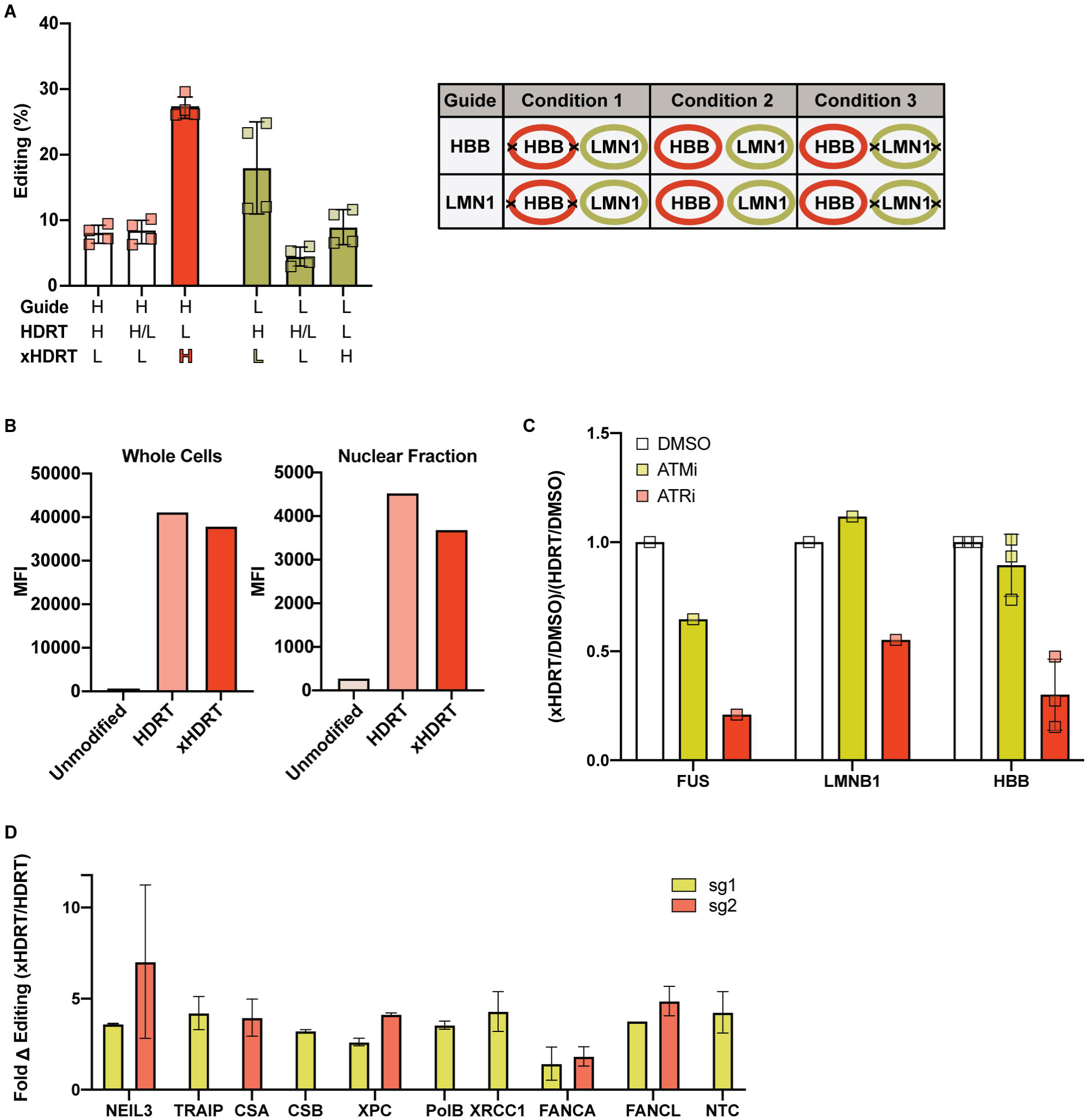
Enhanced editing from xHDRTs requires the activity of DNA repair pathways that are partially distinct from those that support HDR from uncrosslinked plasmids. **(A)** ICLs stimulate HDR in *cis*. Percent incorporation of single or multi-kilobase sequences produced by co-introduced crosslinked (xHDRT) and uncrosslinked (HDRT) DNA in K562 cells. gRNAs and donors target the HBB (H) or LMNB1 (L) loci. Data displayed as the mean ± SD of n=4 biological replicates. **(B)** ICLs do not increase nuclear abundance of xHDRTs. Fluorescence intensity (abundance) of Cy5 labeled HDRT or xHDRT DNA in whole K562 cells (left) or isolated nuclei (right) 24 hours post-electroporation. Data displayed is n=1. **(C)** xHDRT activity is ATR-dependent. Relative change in xHDRT editing during inhibition normalized to the relative change in HDRT editing during inhibition for single (FUS, LMNB1) or multi-kilobase (HBB) sequences in K562 cells treated with DMSO, KU55933 (ATM inhibitor), or AZ20 (ATR inhibitor). Data displayed was calculated from n=3 biological replicates (HBB) or n=1 samples (LMNB1, FUS). See **Fig. S3G** for unnormalized data. **(D)** xHDRT activity is partially dependent on the Fanconi Anemia pathway. Fold change in editing supported by xHDRTs normalized against HDRT editing for the indicated knockdowns. NTC = non-targeting knockdown. Data presented was calculated from at least n=4 biological replicates and multiple independent guides were shown where indicated. Knockdown efficiency is shown in **Fig. S3C**.

We next tested if the xHDRT effect was caused by an increased nuclear abundance of our xHDRTs. Template DNA (crosslinked and uncrosslinked) was nonspecifically labeled with Cy5 and nucleofected into K562 cells. Whole cells or purified nuclei were measured twenty-four hours after nucleofection to recover donor abundance. In both HDRTs and xHDRTs, the majority of the Cy5 signal was associated with the cell membrane or cytoplasm, with no increase in nuclear abundance of xHDRTs (**Fig. 3B**), indicating that ICLs do not increase the nuclear abundance of xHDRTs relative to uncrosslinked templates. It has been reported that biophysical alterations that change the size of RNP particles can improve editing outcomes^8^. We added anionic polymers (ssDNA) to editing reactions containing xHDRTs or uncrosslinked donors and observed robust increases in HDR in all contexts (**Fig S3A**), indicating that xHDRTs act independently from the anionic polymer effect. Together, these results indicate that higher levels of editing seen with xHDRTs requires recognition and processing of the template molecule.

To define these mechanisms, we recovered xHDRT-edited samples into media containing small molecule inhibitors of the apical DNA repair kinases ATM^24^ (5μM KU55933) and ATR^25^ (400nM AZ20), which have both previously been inhibited to alter the frequency and type of DSB repair outcomes^26^. We found that ATR inhibition profoundly (>5-fold) reduces HDR from xHDRTs while modestly (1.3-fold) reducing HDR from uncrosslinked templates. In contrast, ATM inhibition modestly (1.3 fold) reduced the HDR frequency of both uncrosslinked templates and xHDRTs, but xHDRTs still supported higher levels of HDR than uncrosslinked templates **(Figure 3C and S3B)**. ATM (5μM KU55933) or ATR (400nM AZ20) inhibition inhibited the phosphorylation of downstream targets Chk2 and Chk1, indicating that kinase inhibition was effective at these doses (**Figure S3C**). The substrates of ATM and ATR are diverse and overlapping^27^, but the observation that ATR signaling is required for xHDRT repair indicates that uncrosslinked donors and xHDRTs are processed by different DNA repair pathways. Our results are most consistent with a model in which multiple DNA repair pathways can utilize uncrosslinked template DNA, but xHDRTs are strictly processed by ATR-dependent mechanisms.

Due to the local effect of the ICL, we hypothesized that DNA repair factors recruited to the ICL might prime the xHDRT for use as a template. Major pathways implicated in ICL-repair are the Fanconi anemia (FA) pathway, the nucleotide excision repair (NER) pathway, the base-excision repair (BER) pathway, and the NEIL3 glycosylase pathway^19^. To test if these pathways are required for the xHDRT-mediated increase in HDR outcomes, we constitutively inactivated ICL-repair pathways in K562 cells and measured HDR using uncrosslinked and crosslinked template DNA. We separately knocked down genes from each pathway using stably integrated CRISPRi constructs (**Fig S3D**). Knockdown of FANCA significantly attenuated editing from xHDRT relative to uncrosslinked controls (**Fig 3D, S3E**). We also observed attenuation of xHDRT editing in squamous cell carcinoma lines depleted of FANCA (**Fig S3F**). We therefore conclude that the Fanconi Anemia pathway plays a role in increased HDR from xHDRTs.

## Discussion

Our previous work showed that the FA pathway is required for HDR outcomes after Cas9-mediated genome editing, but overexpression of individual FA proteins did not boost HDR frequencies^11^. We therefore investigated if adding ICLs – a substrate of the FA pathway – to donor DNA increased the probability that these molecules would be used for HDR. Strikingly, we found that adding interstrand crosslinks (ICLs) to donor DNA in gene editing reactions dramatically enhances the frequency with which the template is utilized in HDR. This enhancement occurred in many different cell types, and across a range of donors and editing reactions. We also observed that xHDRTs can be used synergistically with other strategies to boost editing efficiency, suggesting a distinct mechanism of HDR enhancement.

We also uncover the outlines of this mechanism: xHDRT editing requires ATR signaling and is partially dependent on the Fanconi Anemia pathway. These dependencies change our understanding of gene editing in several ways. First, the dependence on ATR, which is primarily activated through RPA^28^, suggests that signaling from ATR-activating nuclear structures – and not the DSB – may play a key role in specifying HDR instead of EJ repair pathways. These

ATR-activating structures are unlikely to be encoded on the xHDRT, as these xHDRT molecules do not act as an agonist of ATR (**Figure S3C, ATRi, lanes 1 and 5**), and may instead comprise sites of replication stress or resected DNA. Second, the requirement for ATR in xHDRT editing contrasts with the partial requirement for ATR in uncrosslinked template editing and hints that DNA repair pathway usage may be more heterogeneous than the binary EJ/HDR model (**Fig S1A**) would indicate. One possible explanation for our observations is that uncrosslinked templates can produce HDR outcomes using DNA repair pathways that are ATR-dependent and ATR-independent, while xHDRTs can only be utilized in ATR-dependent pathways. Overall, we favor a model in which xHDRT ICLs are uncovered and repaired during HDR itself and the repair of these lesions, and the completion of HDR, requires ATR signaling.

While our genetic results suggest the Fanconi Anemia pathway is involved in xHDRT processing, the precise mechanism of ICL recognition remains unclear. Proposed mechanisms for FA-mediated ICL repair stipulate that DNA replication uncovers lesions, but degradation rates of HDRTs in cells are inconsistent with episomal replication of these elements **(Figure S3G)**. There are additional models for transcription-coupled repair of ICLs, but these ICL-repair pathways are not required for xHDRT editing (**Fig 3D**). We also note that xHDRTs lacking any eukaryotic promoters support increased levels of HDR **(Figs 2 and S2 LMNB1 and BFP)**. An intriguing possibility is therefore that xHDRT ICLs are uncovered during recombination between the DSB and the template. More work needs to be done to determine if xHDRTs are recognized by known ICL repair pathways, or if a novel recognition mechanism is involved.

From a practical standpoint, xHDRTs support higher levels of HDR with multiple payloads and loci, and in multiple cell types. We thus introduce xHDRTs as a useful tool for laboratory gene editing workflows. Using commercial reagents and the qPCR assay outlined in this manuscript to optimize crosslink density, milligram-scale xHDRT preparations can be completed in a day. Future developments of this approach may enable faster and more effective *ex vivo* cell therapy manufacturing.

## Materials and Methods

### Cell lines and culture

HEK293T and K562 cells were obtained from ATCC, and UMSCC1 cells were obtained from the Fanconi Anemia Research materials repository, held in partnership with the Oregon Health & Science University. K562 cells were cultured in RPMI medium supplemented with 10% fetal bovine serum, 1% sodium pyruvate and 100□μg ml^−1^ penicillin-streptomycin. HEK293T and UMSCC1 cells were cultured in DMEM media supplemented with 10% fetal bovine serum, 1% sodium pyruvate and 100□μg ml^−1^ penicillin-streptomycin. For routine passaging, cells were grown to ∼70% confluency, washed with 1-3 ml DPBS, and subsequently treated with 1-2 ml 0.25% trypsin-EDTA (Gibco) for 3-5 minutes in a 37ºC incubator. Lifted cells were then quenched with DMEM media supplemented with 10% fetal bovine serum, 1% sodium pyruvate and 100□μg ml^−1^ penicillin-streptomycin. Cell lines were routinely tested for mycoplasma contamination using enzymatic (Lonza) and PCR-based assays (Bulldog Bio).

### qPCR-quantification

Purified xHDRT or HDRT plasmids were diluted to 1×10^9^ and 1×10^8^ copies per μl based on measured concentration (Qubit BR kit, Thermo Fisher; or Hoescht 33342). Diluted plasmids were analyzed by qPCR using primers annealing to the ampR gene (oCR3187: cagtgaggcacctatctcagc, oCR3188: taagccctcccgtatcgtagt). Delta Ct values were calculated between the HDRT and xHDRT molecules after pooling of technical triplicates. Delta Cts were averaged between two concentrations of input DNA.

### Cas9, RNA, and HDRT preparation

*S. pyogenes* Cas9-NLS was obtained from the QB3 MacroLab at UC Berkeley.

All sgRNAs were synthesized by Synthego as modified gRNAs with 2’-O-methyl analogs and 3’ phosphorothioate internucleotide linkages at the first three 5’ and 3’ terminal RNA residues.

All dsDNA was derived from purified plasmid DNA from bacterial cultures containing the indicated plasmid (Qiagen Plasmid Plus) or by SPRI purification of amplified linear dsDNA.

Psoralen-mediated xHDRTs were generated by preparing double-strand DNA to a concentration of 100μg/ml in 1X T.E. buffer in a 1.5 ml micro-centrifuge tube. Psoralen (20mM in DMSO) was then added to the reaction tube to the desired final concentration. Each reaction mixture in an open microfuge tube, placed on ice, was then irradiated with long wavelength UV for 15 minutes in a Spectrolinker™ XL-1000. Non-reacted psoralen was removed by an isopropanol precipitation and crosslinked DNA was resuspended in 50μL of 1x TE.

Cisplatin-mediated xHDRTs were generated by diluting double-strand DNA to a concentration of 100μg/ml in 1X T.E. buffer in a 1.5 ml micro-centrifuge tube. Cisplatin (3.3mM in 0.9% saline) was added to the reaction tube to the desired final concentration. The reaction was briefly vortexed and transferred to a 37ºC incubator for one hour. Non-reacted cisplatin was removed by isopropanol precipitation and crosslinked DNA was resuspended in 50μL of 1x TE.

### Cas9 RNP assembly and nucleofection

Per nucleofection, 0.50 μl of sgRNA (100μM) were added to 1 μl of 5x RNP buffer (100 mM HEPES, 750 mM KCl, 25 mM MgCl2, 25% glycerol, 5 mM TCEP) in a 1.5 ml microcentrifuge tube. 1μl of Cas9 protein (40μM) was added to the reaction mixture and then brought up to a volume of 4 μl with nuclease-free water. 1μg of dsDNA donor, prepared at 1 μg/μl, was then added to the RNP mixture. Each reaction mixture was then left to incubate for at least 5 minutes at room temperature to allow RNP formation. 2.5×10^5^ cells were collected and spun down at 500g for 3 minutes, washed once in 200 μl D-PBS, and resuspended in 15μl of nucleofection buffer (Lonza). RNP mixtures were then added to resuspended cell pellets. Reaction mixtures were electroporated in 4D Nucleocuvettes (Lonza), and later recovered to culture dish wells containing pre-warmed media.

Editing was measured at defined time points after electroporation by flow cytometry (standard times are 96 and 120 hours; 240 hours for RAB11A editing – due to transcription off the plasmid). Resuspension buffer and electroporation conditions are as follows for each cell line: K562 in SF with FF-120, UMSCC1 in P3 with DS-138, HEK293T in SF with DS-150, iPSC in P3 with CA-137, T-cell in P3 with EH-115. Viability was measured at defined time points after electroporation by flow cytometry (standard times are 24 and 48 hours).

### Western Blot

∼400,000 cells were lysed in 150μl of 2X Laemmli buffer *(20% glycerol, 120 mM 1M Tris-HCl pH 6*.*8, 4% SDS, 0*.*05% bromophenol blue) containing 100 mM DTT*. Samples were vortexed for 10 seconds at full speed, boiled for 8 minutes, and passed three times through a 25G needle. Whole cell extracts were separated via electrophoresis on Biorad TGX gels 4-20%. Prior to transfer, TGX chemistry was activated for 45 seconds and subsequently used as a loading control. Gels were transferred onto PVDF membranes and blocked for an hour in PBS 0.1% tween and 5% milk. Membranes were incubated overnight in primary antibodies diluted in PBS 0.1% tween and 3% BSA. Membranes were washed in PBS 0.1% tween three times for 10 minutes and incubated for an hour at RT with HRP secondary antibodies (1:5000). Membranes were finally imaged on a Chemidoc (Image Lab™, BioRad). Phospho-Chk1 (1:1000) was detected using antibody #2348 from Cell Signaling. Phospho-Chk2 (1:1000) was detected using #2661 from Cell Signaling. GFP was detected using #A11122 from ThermoFisher (1:2000).

### Dox-inducible transcription

K562 cells stably expressing the reverse tetracycline transactivator (RTTA - Addgene 26429) were nucleofected using a modified LMNB1 donor expressing mCherry under a Tet promoter (PCR 2070). mCherry expression from the donor plasmid was monitored by flow cytometry upon dox induction (1ug/ml).

### T Cell isolation and culture

T cell isolation and culture were performed as previously described^29^. Peripheral blood mononuclear cells (PBMCs) were isolated from fresh healthy donor (donor A) blood by Ficoll centrifugation using SepMate tubes (STEMCELL, per manufacturer’s instructions), or purchased as purified PBMCs (Donor C, STEMCELL). Donor A and C T cells were further isolated from PBMCs via magnetic negative selection using an EasySep Human T Cell Isolation Kit (STEMCELL, per manufacturer’s instructions). Isolated T cells were cultured at 1 million cells ml^−1^ in ImmunoCult medium (STEMCELL) with 5% fetal bovine serum (Bio techné), 50 μM 2-mercaptoethanol (Sigma), and 10 mM N-Acetyl L-Cysteine (Sigma), and were stimulated for two days prior to electroporation with anti-human CD3/CD28 magnetic dynabeads (ThermoFisher) at a beads to cells concentration of 1:1, along with a cytokine cocktail of IL-2 at 200 U ml^−1^ (STEMCELL), IL-7 at 5 ng ml^−1^ (STEMCELL), and IL-15 at ng ml^−1^(STEMCELL). T cells were harvested from their culture vessels and de-beaded on a magnetic rack for several minutes. Prior to nucleofection, de-beaded cells were centrifuged for 3 min at 500g, media was gently aspirated from the pellet, and cells were resuspended in buffer P3 (Lonza), in which 15 μL of buffer were used per one million T-cells.

### T Cell nucleofections

RNPs were made prior to electroporation as described above. One million T cells were de-beaded for several minutes prior to nucleofection and pelleted at 500g for 3 minutes. The cell pellet was then washed with D-PBS. D-PBS was gently aspirated from the T cell pellet and then resuspended in 15 μL of buffer P3 (Lonza). The cell suspension was then transferred to the RNP mix and thoroughly triturated. Next, the cell suspension was transferred to the well of a 20 μL nucleocuvette and immediately nucleofected using the pulse code EH115. Post-nucleofection, cells were rapidly recovered in 1 ml of prewarmed media. Recovery media was composed of ImmunoCult with 5% fetal bovine serum, 50 μM 2-mercaptoethanol, 10 mM N-Acetyl L-Cysteine, and 500 U mL^−1^ IL-2.

### iPSC culture

iPSCs (AICS-0090-391) were acquired from the Allen Institute and treated essentially as described^30^. Low-passage iPSCs were thawed and cultured in 10 ml sterile-filtered mTeSR1 (STEMCELL), without antibiotic, in a 10cm^2^ Matrigel-coated plate and grown to 70% confluency, five days post-thaw. For routine passaging, at 70% confluency, old media was aspirated and cells were washed with 5 ml room temp DPBS prior to dissociation. iPSCs were then treated with 3 ml pre-warmed Accutase (Innovative Cell Technologies) and the vessel was then incubated at 37ºC for 5 min. Once cells began to detach, 3 ml DPBS were added to the Accutase-treated cells and dissociated cells were triturated. Cells were rinsed with an additional 7 ml of DPBS for a final wash, and the dissociated cell suspension was transferred to a 15 ml conical tube and centrifuged at 500g for 3 min at room temp. DPBS/Accutase supernatant was carefully aspirated and cells were resuspended in 10 ml fresh mTeSR1 containing ROCK inhibitor (ROCKi) and counted using a Countess slide. Cells were then seeded into a Matrigel-coated six-well dish at a density of 1.5E + 05 per well in 3 ml mTeSR1 containing ROCKi. Old media containing ROCKi was aspirated from each well the next day and replaced with fresh mTeSR1 *<u>without</u>* ROCKi. mTeSR1 was changed daily, and ROCKi was used for each passaging event, and always removed 24 hours thereafter. All cell line and primary cell work was approved by UCSB BUA2019-15.

### iPSC pre-assembly of Cas9 RNP

For each iPSC nucleofection, 1 μL of 5x RNP buffer (5x stock = 100 mM HEPES, 750 mM KCl, 25 mM MgCl2, 25% glycerol, 5 mM TCEP) and 2 μL of sgRNA (100uM) were mixed with 1.5 μL of 40μM Cas9 protein (QB3 MacroLab) in a microcentrifuge tube along with 1 μg of DNA and brought up to a volume of 6 μL with nuclease-free water. The RNP reaction was incubated at room temperature for 20 minutes.

### iPSC Cas9 RNP Delivery

iPSC RNPs were made prior to electroporation as described above. Low-passage iPSCs, at 70% confluency, in the wells of a six-well Matrigel-coated plate were washed with 2ml D-PBS. D-PBS was aspirated and then 1 ml pre-warmed Accutase was added to each well. Accutase-treated cells were then incubated at 37ºC for 3-5 minutes. 2ml D-PBS were added and lifted cells were triturated, followed by the addition of another 3 ml D-PBS for a final wash. Lifted cells were then transferred to a 15 ml conical tube and pelleted at 500g for 3 minutes. Cells were then resuspended in 10ml fresh mTeSR1 with ROCKi and counted using a Countess slide. 4E + 05 cells were aliquoted per nucleofection and pelleted at 300 g for 5 minutes. Media was aspirated and cells were washed again with D-PBS. D-PBS was aspirated and cells were resuspended in 15 μL buffer P3 (Lonza). The cell suspension was then transferred to the RNP mix and thoroughly triturated in the RNP mix. 20 μL of the resulting cell suspension was carefully, (avoiding the introduction of bubbles), transferred into the well of a 20 μL nucleocuvette (Lonza;). Cells were immediately nucleofected using the ‘Primary Cell P3’ program and ‘CA-137’ pulse code. Post-nucleofection, cells were immediately recovered into the well of a pre-coated 12-well Matrigel plate containing 1 ml of mTeSR1 and ROCK inhibitor. Nucleofected cells were cold-shocked for two days post-nucleofection at 32ºC, transferred to the 37ºC incubator three days post-nucleofection. mTeSR1 media was changed the day after nucleofection, *<u>without</u>* ROCKi. Cells were grown to 80% confluency (typically three days post nucleofection), and passaged using Accutase and ROCKi. Cells were then flowed at 96 hours and 120 hours post-electroporation to measure editing.

### Genomic DNA extraction (for amplicon sequencing)

Approximately 1e+06 cells were harvested two days post nucleofection and incubated in 200 μL of QuickExtract DNA Extraction Solution (Lucigen) at 65ºC for 15 minutes, 68ºC for 15 minutes, and 95ºC for 15 minutes. Extracts were diluted 1:4 with dH_2_O and insoluble cell debris was removed by centrifugation. Supernatants were then transferred to a new tube for downstream analysis.

### PCR amplification of edited regions

Edited loci were amplified using locus-specific primer pairs described in Supplementary Table 1 using GoTaq master mix (Promega) and 200ng of genomic DNA. The thermocycler was set for 1 cycle of 98°C for 30s, 35 cycles of 98°C for 10s, 62°C for 10s and 72 °C for 30 s, respectively, and 1 cycle of 72 °C for 1 minute. PCR amplicons (PCR1) were purified using SPRI beads, run on a 1.0% agarose gel to validate size and quantified by Qubit. 100ng of purified PCR1 DNA was then reamplified with PCR2 primers as listed in Supplementary Table 1. PCR conditions are in order as follows, 95ºC for 2 minutes, 95ºC for 30s, 60ºC for 20 cycles, 72ºC for 30s, 72ºC for 2 minutes. PCR2 products were SPRI cleaned, quantified by Qubit, normalized, and pooled at equimolar amounts. PCR2 pools were sequenced using 2×300 chemistry on a Miseq.

### Analysis of Amplicon Sequencing Data

Reads were adaptor and quality trimmed using trim_galore (version 0.6.6) and aligned to predicted amplicon sequences using bowtie2 (version 2.2.5, --very-sensitive-local mode). Nucleotide variants at each position of the aligned reads were quantified using bcftools mpileup and bcftools call (version 1.11-1-g87d355e, m -A flags passed to bcftools call). Nucleotide variants were extracted using bcftools query in two formats: all nucleotides in a 50bp window centered on the cut site, and all nucleotides in a 50bp window centered on the cut site with HDR nucleotides removed. These values were plotted on a per-nucleotide basis [**Fig S2C**] or summed to produce bar plots [**Fig S2B**].

### Nuclear localization experiments

HDRT and xHDRT DNA were Cy5-labeled using the *Label* IT® Nucleic Acid Labeling Reagents (Mirus) and used in a standard nucleofection protocol (see Cas9, RNP assembly and nucleofection, with about 1×10^6^ cells). At 6 and 24 hours, 5×10^5^ cells were collected and washed in PBS. 10% of the cells were analyzed by flow cytometry (WCE fluorescence). The rest of the samples were processed for nuclei isolation as follows: cells were resuspended in 475 μl of hypotonic buffer (20mM Tris-HCl, pH 7.4, 10mM NaCl, 3mM MgCl_2_) and incubated on ice for 15 minutes. 25μl of 10% NP40 were added and the samples were vortexed full speed for 20 seconds. Nuclei were spun for 5 minutes at 700g and resuspended in PBS. Nuclei were then assessed by flow cytometry. The quality of the nuclei was ascertained by analyzing the FCS/SSC channels (nuclei should be about a third of the size of the whole cell extract).

### ATR/ATM inhibition experiments

After standard nucleofection, cells were recovered in media containing the indicated concentration of ATR inhibitor (AZ20) or ATM inhibitor (KU55933).

### Lentiviral packaging

Lentiviral packaging was adapted from^31^. Lentivirus was produced by transfecting HEK293T cells with standard packaging vectors using the TransIT-LT1 Transfection Reagent (MIR 2306; Mirus Bio LLC). Viral supernatant was collected 48–72 h after transfection, snap-frozen and stored at −80 °C for future use.

### CRISPRi knockdown

Plasmids encoding gRNAs targeting FANCA, FANCL, NEIL3, XPC, CSA, CSB, DDB1, POLB, TRAIP, XRCC1, or a non-targeting sequence (Supplementary Table 1) were separately transduced into K562 cells containing dCas9-KRAB (clone K1e [Richardson et al Nat Gen 2017]). Resulting cell populations were selected to homogeneity using puromycin (1μg/mL). Pooled knockdown cell populations were tested as described in the manuscript, and knockdowns validated by qPCR.

### qPCR for CRISPRi cell-lines

For qPCR, 2.5×10^5^-1×10^6^ CRISPRi cells were harvested. RNA was extracted using RNeasy Mini Kits (Qiagen). RNA was quantified by nanodrop and cDNA was produced from 1□μg of purified RNA using the iScript Reverse Transcription Supermix for RT-qPCR (Bio-Rad Laboratories). qPCR reactions were performed using the SsoFast Universal SYBR Green Supermix (Bio-Rad Laboratories) in a total volume of 10□μl with primers at final concentrations of 500□nM. The thermocycler was set for 1 cycle of 95□°C for 2□min, and 40 cycles of 95□°C for 2□s and 55□°C for 8□s, respectively. Fold enrichment of the assayed genes over the housekeeping control *ACT1B* locus were calculated using the 2^− ΔΔ*C*^T method essentially as described.

### Cell Cycle Experiments

Cell cycle analysis was performed using Click-iT EdU Alexa Fluor 647 Flow Cytometry Assay Kit (#Thermofisher C10424) with the following modifications: Cells were pulse labelled with EdU at 10uM final for 30min, fixed in 4% formaldehyde for 10min, washed twice with PBS containing 1% PSA, permeabilized for 15min with PBS containing 0.5% triton X100. Click iT reaction was carried out following manufacturer instruction. After 3 washes with PBS containing 1% BSA, cells were treated for 30min with RnaseA, and stained for 10min with propidium iodide and run on the flow cytometer.

### Data Availability

Data from amplicon sequencing will be available upon publication at [location].

## Figure Legends

**Figure S1:**
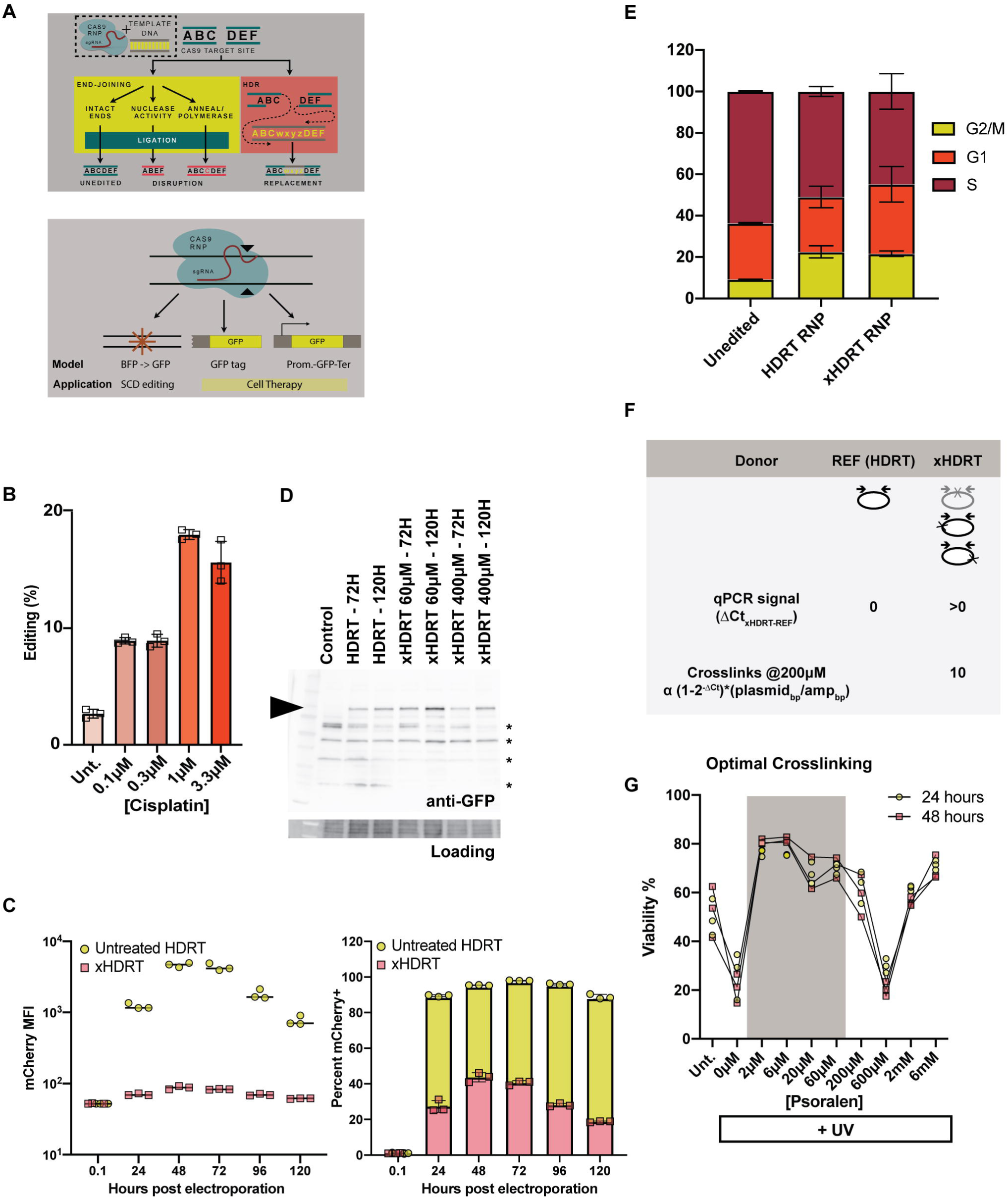
Modification of HDRTs with interstrand crosslinks increases HR during gene editing. **(A)** Top panel: Cas9 RNPs introduce a double strand DNA break (DSB) at a targeted region in the genome, which can be repaired by error prone end joining (EJ) processes that rejoin the ends of the break, or homology-directed repair (HDR) processes that resolve DSBs using sequence encoded in a separate template molecule. Bottom panel: HDR gene editing applications include SNP replacement and single or multi-kb sequence insertion and can be approximated using marker-based assays as diagrammed. These editing events are initiated by electroporation of Cas9, sgRNA, and HDRT into human cells, and monitored by flow cytometry or high throughput sequencing. **(B)** Incorporation frequency of a pSFFV-GFP construct into the HBB locus of K562 cells using plasmid DNA treated with the indicated amount of cisplatin. **(C)** Transcription is inhibited from xHDRTs. Expression of dox-inducible mCherry presented both as intensity (left) of uncrosslinked and xHDRT DNA and percent of cells expressing mCherry (right). Data displayed as the mean ± SD of n=3 biological replicates. **(D)** Western blot for GFP in K562 cells edited with xHDRTs that insert GFP at the N-terminus of LMNB1. Cross-reacting bands (asterisks) are shown on the blot. Blot is representative of n=3 biological replicates. **(E)** xHDRTs do not alter the cell cycle more than uncrosslinked templates. Percent of asynchronous cells edited with uncrosslinked donors or xHDRTs at the indicated point in the cell cycle. Data displayed as the mean ± SD of n=3 biological replicates **(F)** Schematic of crosslinking quantification by qPCR. Untreated (HDRT) or xHDRT molecules were amplified using PCR primers that produce a 94bp amplicon. Cycle thresholds (Cts) were calculated for each sample and subtracted from an uncrosslinked control to obtain ΔCt. ΔCt numbers were converted back to the number of crosslinks using the indicated formula. We calculate 10 crosslinks per kilobase for the 60μM xHDRTs. **(G)** Viability of human K562 cells 24 or 48 hours after editing using xHDRTs with varying numbers of crosslinks.

**Figure S2:**
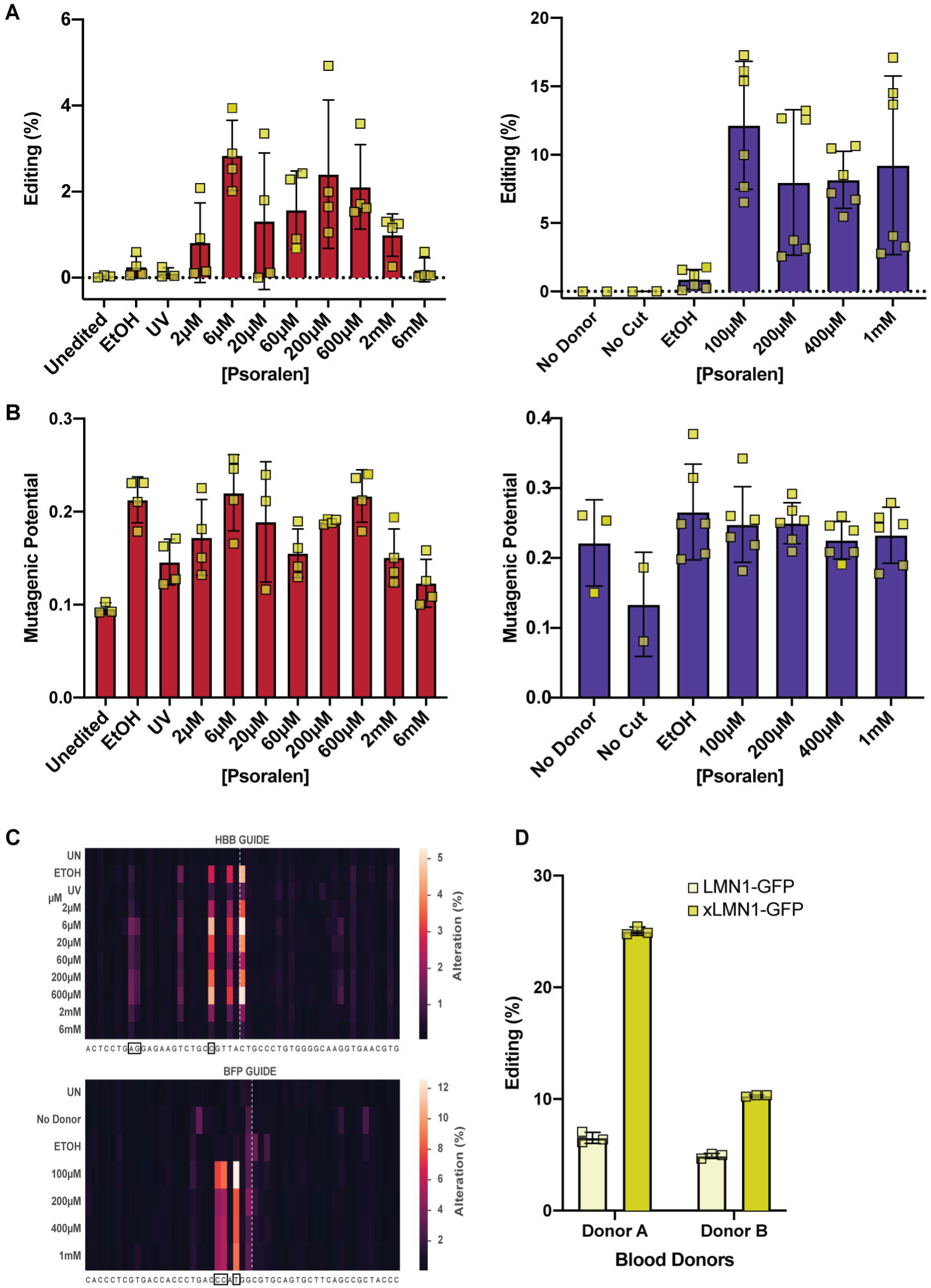
xHDRTs increase HDR in broad gene editing applications. **(A)** xHDRTs boost SNP conversion. SNP conversion as a function of crosslink frequency at the HBB (left) or BFP (right) loci in K562 cells. Data displayed as the mean ± SD of n=4 biological replicates. **(B)** xHDRTs do not increase total number of mutations in a window surrounding the cut site. Cumulative probability of a non-HDR mutation (mutagenic potential) arising within a 50bp window surrounding the Cas9 cut site for samples edited with the indicated homology donors at HBB (left) or BFP (right) loci in K562 cells. **(C)** xHDRTs do not increase the mutation frequency at non-SNP bases. Heatmap showing mutation frequency at each base within a window surrounding Cas9 cut site (white dashed line) for samples edited with the indicated homology donors at HBB or BFP. Nucleotides altered by successful HDR are outlined with black squares. **(D)** xHDRTs enhance HDR in primary T-cells. Percent incorporation of a single-kilobase construct at the LMNB1 locus in primary T-cells from two donors. Data shown are the mean ± SD of n=3 biological replicates.

**Figure S3:**
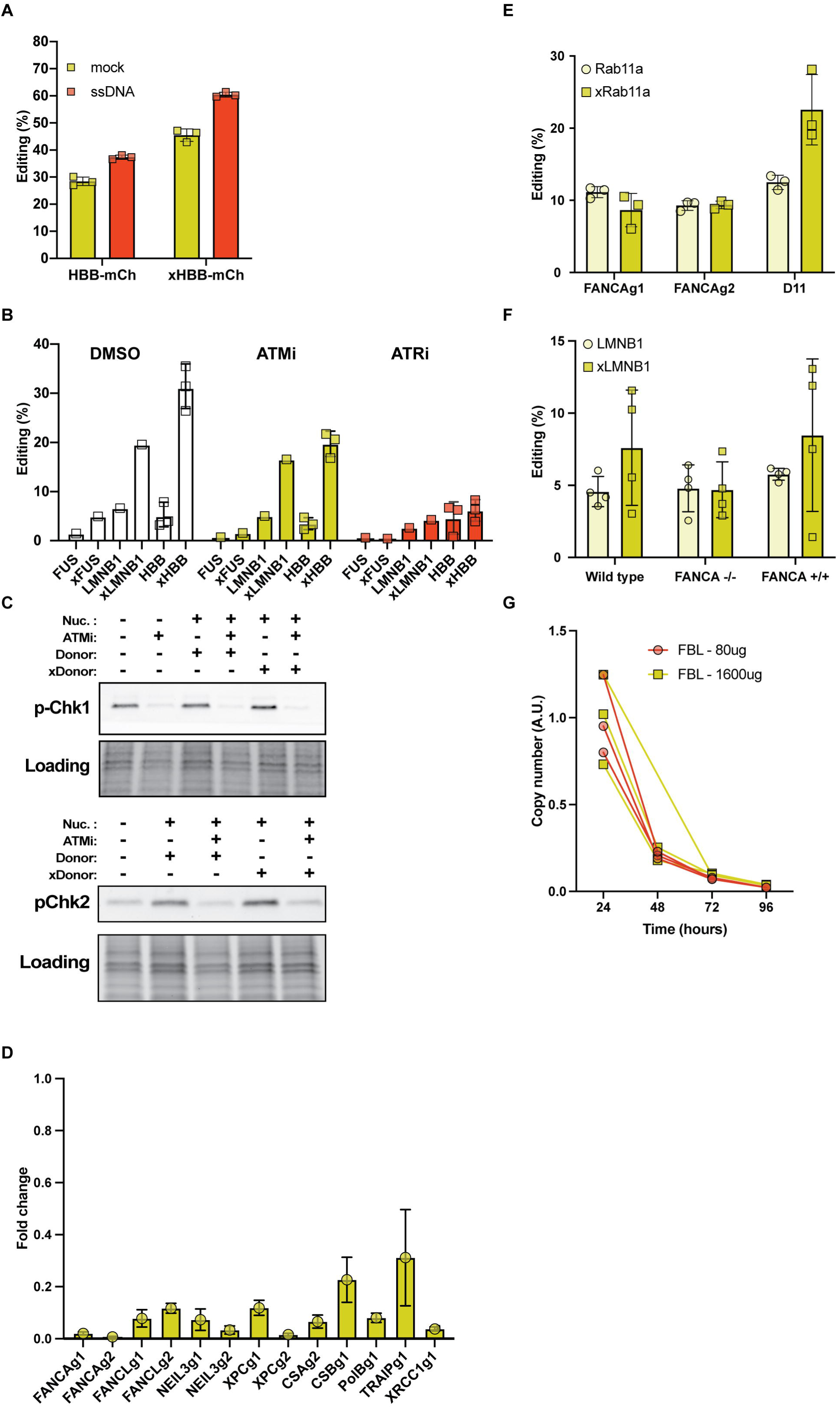
Enhanced editing from xHDRTs requires the activity of DNA repair pathways that are partially distinct from those that support HDR from uncrosslinked plasmids. **(A)** xHDRTs work additively with the anionic polymer effect. Percent incorporation of a multi-kilobase (HBB-mCherry) construct with or without 100 pmoles of nonhomologous ssDNA. Data shown are the mean ± SD of n=3 biological replicates. **(B)** xHDRT activity is ATR-dependent. Percent incorporation of single (FUS, LMNB1) or multi-kilobase (HBB) sequences using crosslinked (xHDRT) and uncrosslinked DNA in K562 cells treated with DMSO, KU55933 (ATM inhibitor), or AZ20 (ATR inhibitor). Data displayed as the mean ± SD of n=3 biological replicates (HBB) or n=1 samples (LMNB1, FUS). **(C)** ATM and ATR inhibitors prevent substrate phosphorylation. Western blots for phospho-Chk1 (top) and phospho-Chk2 (bottom) 24 hours after the indicated treatments. Data shown is representative of n=3 blots. **(D)** CRISPRi knockdown of ICL-repair genes was effective. Percent knockdown of indicated guides in CRISPRi cell lines as measured by qPCR. Data displayed as the mean ± SD of n=3 biological replicates. **(E)** xHDRT activity is partially dependent on FANCA. Percent incorporation of a single kilobase construct in two independent FANCA knockdown K562 cell lines. Data displayed as the mean ± SD of n=3 biological replicates. **(F)** xHDRT activity is partially dependent on FANCA. Percent incorporation of a single kilobase construct in UMSCC1 cells with the indicated genotype. Data displayed as the mean ± SD of n=4 biological replicates. **(G)** Abundance of donor plasmids decreases over time in cells. qPCR plasmid quantification (AU= 2^(Ct^plasmid^-Ct^genome^)_tN_/2^(Ct^plasmid^-Ct^genome^)_t24_) at the indicated times after electroporation. Half-life is approximately 12 hours.

## Acknowledgments

We thank the Fanconi Anemia Research Fund for enabling the research reported here by providing the UMSCC1 cells from the Fanconi Anemia Research Materials repository that is held in partnership with the Oregon Health & Science University. The authors acknowledge the assistance of Dr. Jennifer Smith (manager of the BNL) and the use of the Biological Nanostructures Laboratory within the California NanoSystems Institute, supported by the University of California, Santa Barbara and the University of California, Office of the President. We thank David N. Nguyen for assistance with T-cell protocols. We thank Cassidy Arnold for assistance with iPSC protocols. Research reported in this publication was supported by the National Institute of General Medical Sciences of the National Institutes of Health under Award Number R35GM142975. The content is solely the responsibility of the authors and does not necessarily represent the official views of the National Institutes of Health.

